# Assessment of Carbon Sequestration Potential of some Selected Urban Tree Species

**DOI:** 10.1101/2021.08.19.457022

**Authors:** Ezekiel Ajayi

## Abstract

Tree species carbon assessment and quantification remain the only opportunity to determine the position of forest in climate change amelioration potentials. Forest biomass constitutes the largest terrestrial carbon sink and accounts for approximately 90% of all living terrestrial biomass. The aim of this study is to assess tree species carbon sequestration potentials of selected urban tree species. The study was carried out in Adekunle Ajasin University Campus, Akungba Akoko, Ondo State, Nigeria. All trees species ≥10 cm Diameter at Breast Height (Dbh) within the area were identified and their Dbh measured as well as other variables for volume computation such as height, diameters at the base, middle and top. Also, for density assessment; stem core samples were collected. Again, the coordinate of individual tree was recorded using a Global Positioning System (GPS) receiver. A total of 124 individual trees were encountered with varying growth variables as well as carbon values. The study area contains some indigenous and exotic tree species such as *Acacia auriculiformis, Terminalia mantily, Gmelina arborea* and *Tectona grandis* etc. but *Acacia auriculiformis* had the highest frequency. The tree species with highest carbon sequestration was *Gmelina arborea* as recorded for this study. The total carbon and carbon dioxide sequestered in the study area were reported as 47.94 kg and 176.03 kg respectively.

## INTRODUCTION

Carbon emission and reflection is one of the most important and pressing environmental issues due to climate change in the world today ^1^. Expansions of urban areas are steadily throughout the entire world ^2^ and it is expected that about sixty percent (60%) of the world population will inhabit in cities by 2030 ^3^. Unfortunately, urbanization will therefore increase the source of carbon emissions in the world. As a result, urban trees become more important for human to enjoy their living space.

Several studies around the world such as North America, China, and Australia as well as United Kingdom and Germany recently had reported that trees in urban environments can remove carbon dioxide from the atmosphere through photosynthesis which results in tree growth in both directions (horizontal and vertical) ^4, 5^. This accumulation of resources stores excess as biomass/carbon in various components of the tree such as roots stems and branches etc. Incidentally, urban trees reduce building energy due to cooling through their shade and climate amelioration effects, thereby reducing carbon dioxide emissions ^6^. It is noteworthy that the strength of urban tree carbon sequestration depends on different factors such as species, health and growth variables as well as their overall living conditions ^7, 8^. Also, the mortality or longevity of urban trees can be influenced by site condition(s) and other policies such as land use, natural disturbance (e.g. pests, fire and desert encroachment, etc.), as well as human activities and urban development ^8^. Comparably, urban tree growth is influenced by genetics make up, climate and competition amongst other ^9^.

Trees/forest in the urban centres provides several ecosystem services ranging from atmospheric cooling to carbon emission ^10^. Although, these importance has not been documented extensively but European Union Biodiversity Strategy 2014 mandated her member states to asses and map their forest ecosystem services in the year 2020 ^11^. Some European cities have begun to formulate carbon dioxide mitigation policies such as; city of Bolzano and Italy. These cities have been designated to become carbon monoxide neutral by 2030 ^12^. Consequently, many countries are on the quest to assess their forest carbon emission and sequestration by using developed allometric equations both regionally or local as well as other forest inventory methods to provide reliable, consistent and scientifically proven forest biomass/carbon estimate for reporting. The assessment of carbon in tree or forest ecosystem gives an estimate of carbon emitted into the atmosphere when this particular tree/forest is deforested or harvested. Consequently, the aim of this study is to assess carbon sequestration potentials of some selected urban tree species.

### Study Area

This study was conducted in Adekunle Ajasin University Campus, Akungba Akoko, Ondo State, Nigeria. It is situated in Akoko South West Local Government Area of Ondo State and lies between latitudes 7^°^ 28” 9.15’ to 7^°^ 29” 15.18’ North of the equator and longitudes 5^°^ 44” 15.96’ to 5^°^ 46” 14.78’ East of Greenwich Meridian. Akungba town is bordered with Supare-Akoko in the West, Iwaro-Oka in the East, Ikare-Akoko in the North and in the South with Oba-Akoko ^13^.

The study area falls in rainforest ecological zone but gradually becoming derived savannah due to anthropogenic activities causing climate change effects ^13^. The area is characterized by two distinct seasons; the raining season, which occur between April and September and the dry season, which falls between September and March. It has mean annual rainfall of 1250 mm and the average temperature ranges between 18°C and 35°C.

## Methods

Stem of living trees grows both horizontally and vertically. Biomass accumulation also occurs in trees in both directions. The horizontal growth was measured by the diameters and the vertical growth was measured by the total tree height. Newton’s formula was used to estimate the total tree volumes for this study (Equation 1).

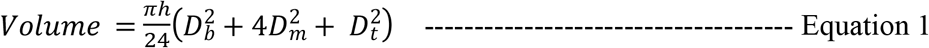

Where:

Volume = Volume of the stem

∏ = 3.142

h = Tree total Height

D_b_ = Diameter at the base

D_m_ = Diameter at the middle

D_t_ = Diameter at the top.

### Density

A non-destructive sampling method was adopted in this present study to estimate the stem biomass. The length of the stem core extracted using the increment borer was accurately measured in centimeter using ruler. Core diameter was measured for randomly selected core samples and the average taken since only one increment borer with one extraction tube was used for the study. The core samples obtained were oven-dried at 75^°^C and measured at one (1) hour interval until a constant weight was achieved. The density estimation was therefore done by converting the volume and dry weight of a core sample extracted to the stem density. Both Equations 2 and 3 were used to estimate tree core volume and tree density respectively.

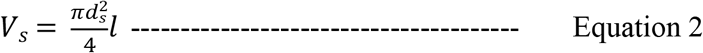

Where;

*d*_s_*=* diameter of the core sample (cm)

*l =* core sample length (cm)

*V*_*s*_*=* volume of the core sample (m)

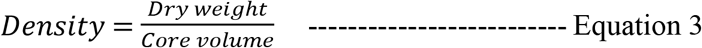

Where:

Density = Tree density

Dry weight = Core sample dry weight

Core volume = Core sample volume

### Estimation of biomass

Biomass of each tree was estimated using the volume and density as estimated from respective tree and Equation 4 was employed to estimate bionmass.

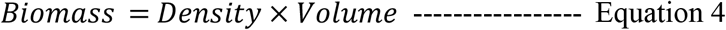

Where:

Density = value obtained in equation 3

Volume = value obtained in equation 1

### Estimation of Carbon

Tree biomass obtained in equation 4 was used to estimate carbon stock for each tree. The standard multiple factor of 0.5 was used for conversion of biomass to carbon (Equation 5) ^14^.

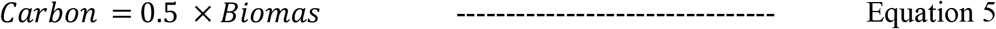

Where:

Biomass = value obtained in equation 4

### Estimation of Carbon dioxide

To convert carbon to carbon dioxide, the carbon values are multiply by the ratio of the molecular weight of carbon dioxide to that of carbon (44/12) ^15^. The value(s) obtained in equation 5 was used to convert carbon to carbon dioxide (CO_2_) using Equation 6.

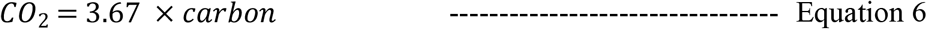

Where:

Carbon = value obtained in equation 5

## Results and Discussions

The result of this study showed 124 individual trees in 18 tree species. The tree species with highest frequency is *Acacia auriculiformis* with 31 individual followed by *Terminalia mantaly* with 22 individual and some species was encountered once e.g. *Ficus exasperata, Hura crespatans, Pilliostigma thonningii, Pinus caribaea, Vitex doniana*, etc.

The highest carbon and carbon dioxide was recorded from *Gmelina arborea* with 25.41kg carbons and 93.03kg carbon dioxide respectively followed by *Acacia auriculiformis* with carbon and carbon dioxide of 4.65kg and 17.05kg respectively. The lowest carbon and carbon dioxide was recorded for *Piliostigma thonningii* with 0.04kg and 0.17kg respectively (Table 1). This result of the carbon and carbon dioxide of *Gmelina arborea from* 21 individual trees is consistent with the result of ^16^ who reported 262.6kg of carbon from 212 individual trees. The total carbon sequestered by *Acacia auriculiformis* (4.65kg), *Albizia lebbeck* (0.63kg), *Tectona grandis* (2.28kg) reported in this study was lower than the values reported by ^17^ which are: 22.891kg for *Acacia auriculiformis*, 15.169kg for *Albizia lebbeck* and 26.071kg for *Tectona grandis* in a research conducted in Jadavpur University, Kolkata, Indian. This may be due to ecological and characteristic of tree as well as condition of tree stocks. The trees waypoint (coordinate) is presented in the appendix 1.

**Table 1:**
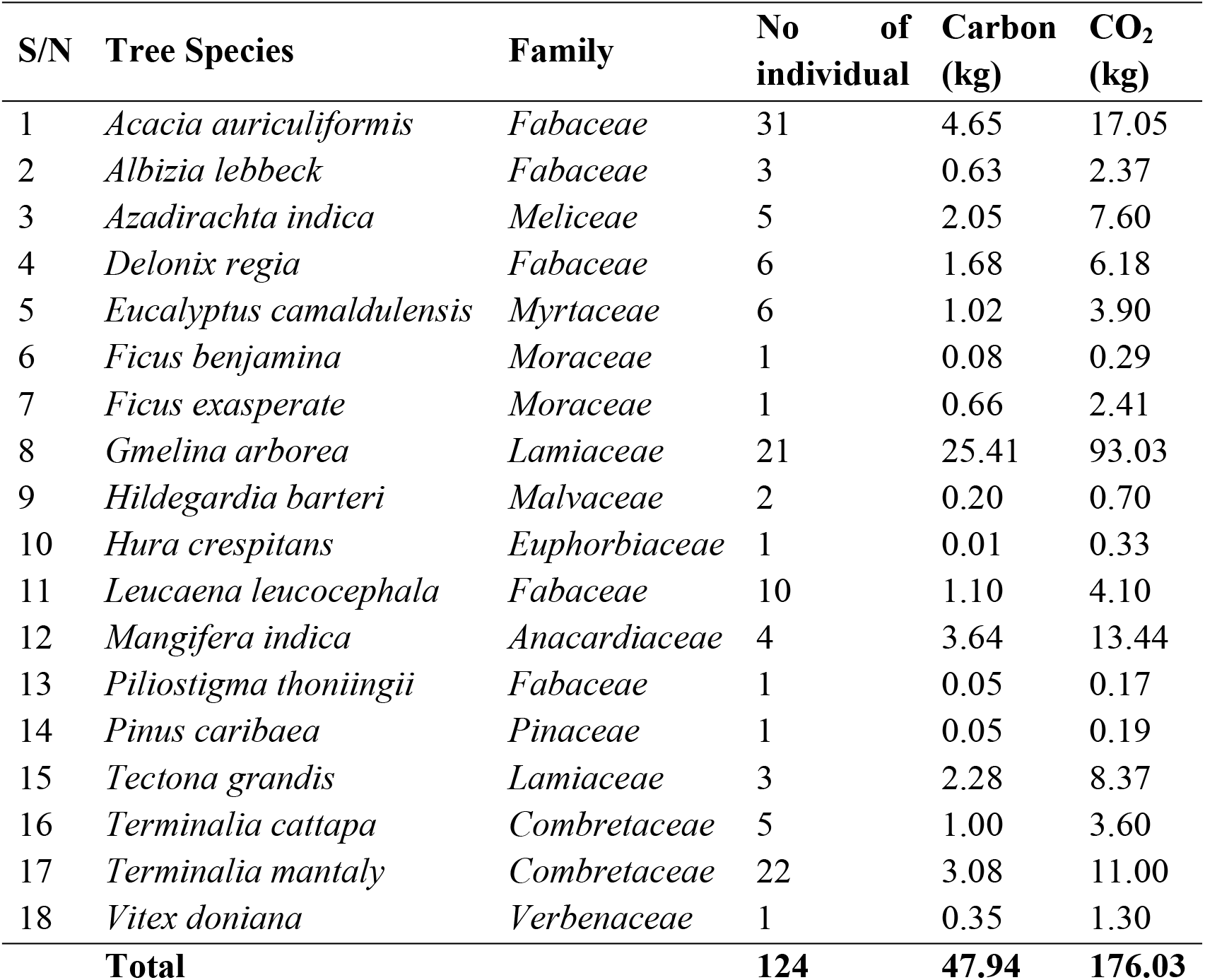
Summary of Species, Family and their Carbon Sequestered

## Conclusions and Recommendations

Basically, every tree generates a different amount of carbon sequestration based on the tree characteristics including tree diameter and tree height. The total amount of carbon and carbon dioxide sequestered by the 124 trees encountered was 47.94kg and 176.03kg respectively. This study had equipped forester as well as urban planner the types of tree with high sequestered carbon and this will in turn inform the choice of tree to be planted in urban area. These species (*Gmelina arborea, Acacia auriculiformis, and Mangifera indica)* could be recommended for planting in any University campus for better sequestration and assimilation of carbon to enrich the quality of ecosystem services within campus community.

## Apendix 1

**Table 2:**
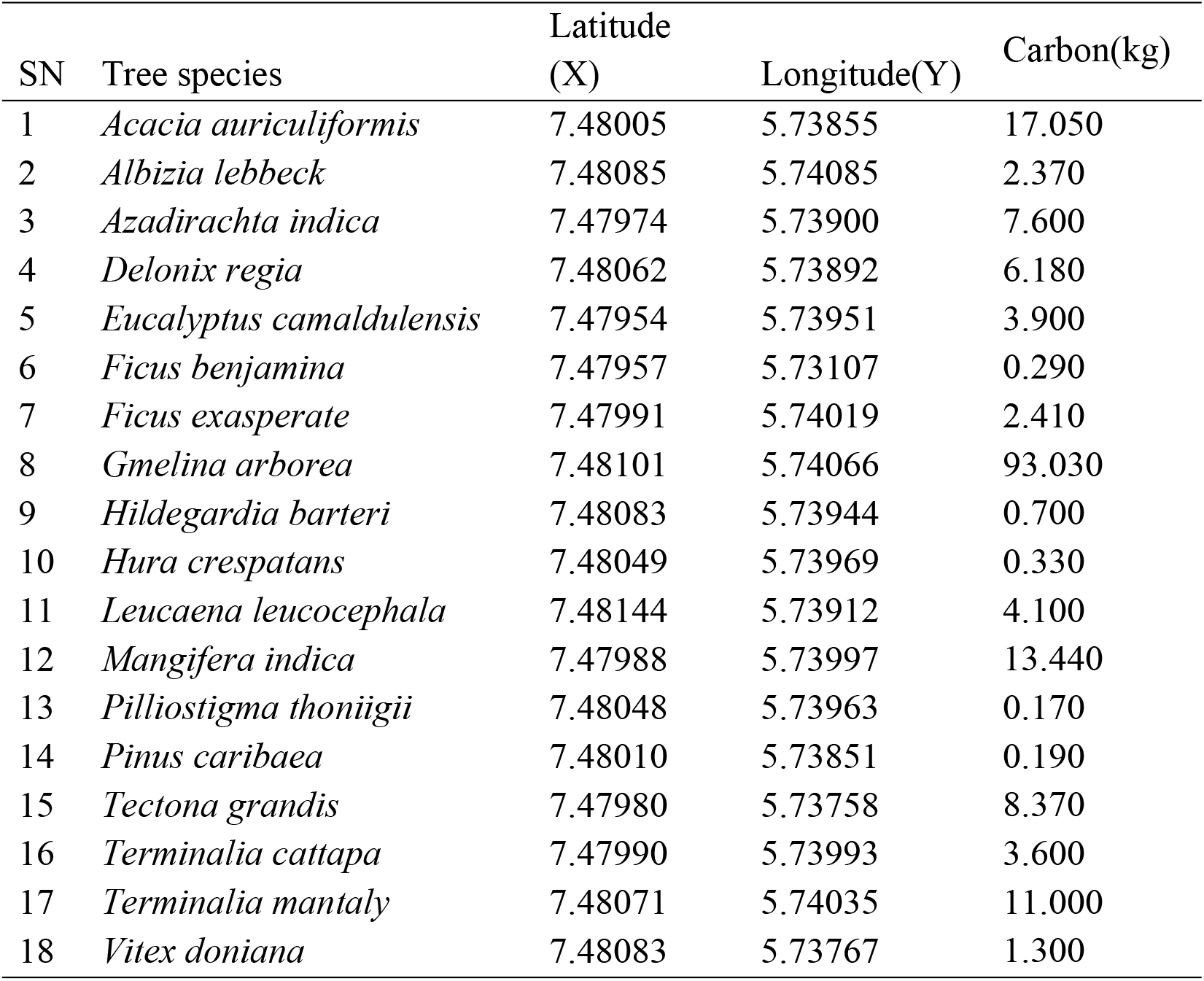
Coordinates of individual tree species and carbon sequestered

